# MDITRE: scalable and interpretable machine learning for predicting host status from temporal microbiome dynamics

**DOI:** 10.1101/2021.12.15.472835

**Authors:** Venkata Suhas Maringanti, Vanni Bucci, Georg K. Gerber

**Affiliations:** Department of Computer and Information Science, University of Massachusetts Dartmouth, MA, USA; Department of Microbiology and Physiological Systems, University of Massachusetts Medical School, Worcester, MA, USA; Program in Microbiome Dynamics, University of Massachusetts Medical School, Worcester, MA, USA; Department of Pathology, Brigham and Women’s Hospital, Boston, MA, USA; Harvard Medical School, Boston, MA, USA; MIT-Harvard Health Sciences and Technology, Cambridge, MA, USA

## Abstract

Longitudinal microbiome datasets are being generated with increasing regularity, and there is broad recognition that these studies are critical for unlocking the mechanisms through which the microbiome impacts human health and disease. Yet, there is a dearth of computational tools for analyzing microbiome time-series data. To address this gap, we developed an open-source software package, MDITRE, which implements a new highly efficient method leveraging deep-learning technologies to derive human-interpretable rules that predict host status from longitudinal microbiome data. Using semi-synthetic and a large compendium of publicly available 16S rRNA amplicon and metagenomics sequencing datasets, we demonstrate that in almost all cases, MDITRE performs on par or better than popular uninterpretable machine learning methods, and orders-of-magnitude faster than the prior interpretable technique. MDITRE also provides a graphical user interface, which we show through use cases can readily derive biologically meaningful interpretations linking patterns of microbiome changes over time with host phenotypes.

## Introduction

The human microbiome is highly temporally dynamic^1^. Some of the most profound changes over time occur during infancy and early childhood when the microbiome is first becoming established^2–4^. Although the microbiome is more stable in adulthood, it continues to undergo significant changes over time due to diet^5,6^, travel^7^, antibiotic use^8^, infection^7^, gut inflammation^9^, and a variety of other factors. Microbial dynamics, particularly early in life, have been linked to many human diseases including necrotizing enterocolitis^10^, diabetes^4,11^, food allergies^12^, obesity^13^, and inflammatory bowel diseases^9^. An increasing number of prospective longitudinal studies have been undertaken to characterize microbiome-disease relationships. Such longitudinal studies are particularly important for advancing the field, as they can aid in establishing causality (e.g., changes definitively proceeding disease onset) and provide information for clinically useful diagnostic or prognostic tests.

Relatively few computational or statistical methods have been specifically developed to analyze longitudinal microbiome data, despite its importance to the field. Human microbiome time-series data present numerous challenges, including small numbers of subjects, high subject-to-subject variability, case/control imbalance, irregular/sparse temporal sampling, high-dimensionality, compositionality, and complex dependencies among variables^1^. Methods that have been developed for analyzing microbiome time-series data generally fall into four categories: (1) univariate models of taxa trajectories (for example, ^14^), which are useful for interpolating data or characterizing differences over time between two cohorts on a taxon-by-taxon basis, (2) dynamical systems models that capture microbe-microbe interactions (for example, ^15,16^), which are useful for forecasting ecosystem behaviors over time such as responses to perturbations or stability, (3) unsupervised learning or clustering methods, which are useful for characterizing common patterns of change among microbes (for example, ^5^), and (4) supervised learning methods that predict host status or outcomes using trajectories of multiple taxa as inputs (for example,^17,18^), which are useful for establishing associations between microbiome dynamics and host phenotypes or developing diagnostic/prognostic tests. Our present work falls into this latter category.

In the supervised learning domain, general-purpose “black-box” machine learning methods, such as deep neural networks and random forests, have become increasingly popular. Indeed, such methods have been applied to microbiome data, and have been demonstrated to predict host phenotype accurately^19,20^, including from longitudinal data^18^. Although these methods can achieve high predictive performance, by their nature they encode mathematical functions that are incomprehensible to humans. In some domains, such as speech recognition for consumer applications, human comprehension of the underlying model is not important. However, in many biomedical domains, including the microbiome, understanding the underlying model is critical; the end-consumers of analyses are often wet-lab experimentalists or clinicians, who ultimately seek to generate specific testable hypotheses or to develop diagnostic/prognostic clinical tests. One approach for understanding black-box models is *post hoc* techniques that attempt to explain individual components of black-box models with simpler models. As an example in the microbiome domain, the Local Interpretable Model Agnostic Explanations (LIME)^21^ technique has been applied to random forests to attempt to find abundance thresholds of specific microbes that differentiate patients according to disease severity^22^. Although these *post hoc* techniques are useful, they suffer from several limitations, including inherent unfaithfulness to the original model or difficulty expressing how inputs are jointly related to each other^23,24^.

An alternative to black-box machine learning methods are models that are purposefully constructed to be interpretable. The notion of interpretability is inherently domain-specific and ultimately hinges on the ability of human experts to comprehend the models^23^. In prior work, we introduced the Microbiome Interpretable Temporal Rule Engine (MITRE)^17^, a fully Bayesian, microbiome time-series specific model that learns human-interpretable rules to classify the host’s status (e.g., healthy, or diseased) from microbiome time-series data. MITRE rules consist of conjunctions of *detectors* that handle dependencies in both microbiome and time-series data. These detectors are conditional clauses of the form: “*TRUE if the aggregated abundance (or rate of change of abundances) of microbes in phylogenetic subtree A within time window T is above threshold B*.*”* This approach, which performs a set of nonlinear but interpretable and domain-specific transformations on the inputs, was demonstrated to perform on par and or outperform black-box machine learning methods (i.e., Random Forests). However, because MITRE uses a sampling-based inference approach, which operates combinatorically on a large space of pre-computed features, it is not scalable to increasingly larger microbiome datasets.

An exciting recent direction is to leverage advances in computing technologies originally developed for black-box deep learning, to greatly accelerate interpretable logic or rule-based approaches. At the core of the deep learning revolution are software and hardware advances, including Graphical Processing Units (GPUs) that perform highly parallelized computations to optimize nonlinear functions using gradient-descent-based methods. These approaches require that the functions to be optimized are differentiable. Standard logic or rule-based models are not differentiable, because logical clauses and their combinations are discrete, not continuous, entities. To tackle this problem, relaxation approaches can be used, which construct smooth approximations to the underlying logical functions that are successively made sharper throughout the learning algorithm^25–27^.

Building on this work, to achieve scalability on large microbiome datasets while maintaining model interpretability, we developed the Microbiome Differentiable Interpretable Temporal Rule Engine (MDITRE), a fully differentiable version of our original MITRE method. The remainder of this manuscript is organized as follows. First, we introduce the MDITRE model, including domain-specific microbiome and temporal focus mechanisms, which enable model differentiability. We also provide details on the MDITRE open-source software package, which can run on GPUs and provides a graphical user interface. Next, we present predictive performance and run-time benchmarking results of MDITRE against MITRE and other methods, on both semi-synthetic and real microbiome datasets, which study a variety of host phenotypes/outcomes. We also show that MDITRE can scale to much larger datasets than MITRE can feasibly run on, through both semi-synthetic and real data. Finally, we provide cases studies illustrating MDITRE’s ability to readily uncover biologically interpretable patterns in datasets, using its automatically inferred rules and visualization capabilities.

## Results

### MDITRE has a fully differentiable architecture that enables scalability while maintaining interpretability

MDITRE is a highly scalable approximation to our method, MITRE^17^, a fully Bayesian supervised machine learning framework that classifies hosts according to specified binary labels (e.g., disease or healthy) using microbiome time-series data. MDITRE takes as input (**Figure 1A**): (1) a binary (two-value) label of the status of each host, (2) a table of microbial longitudinal relative abundances (OTUs or ASVs from 16S rRNA sequencing, or taxa derived from shotgun metagenomic data) and, (3) a table of pairwise distances (matrix of phylogenetic distances among taxa). Using these data as inputs, MDITRE learns human-interpretable *rules*, which explicitly incorporate microbiome- and temporally-specific features, to output predictions of host labels (**Figure 1B,E**). A rule consists of a conjunction (logical AND) of *detectors*. Detectors are of the form “*TRUE if the [aggregated abundance / rate of change of abundance] of taxa in group A within time window T is above threshold Y*.” The label for each host (e.g., healthy or disease) is then predicted by the model based on a weighted sum of rules; the weights on the rules can be interpreted as the odds of predicting a particular host label, given the input microbiome time-series information.

**Figure 1:**
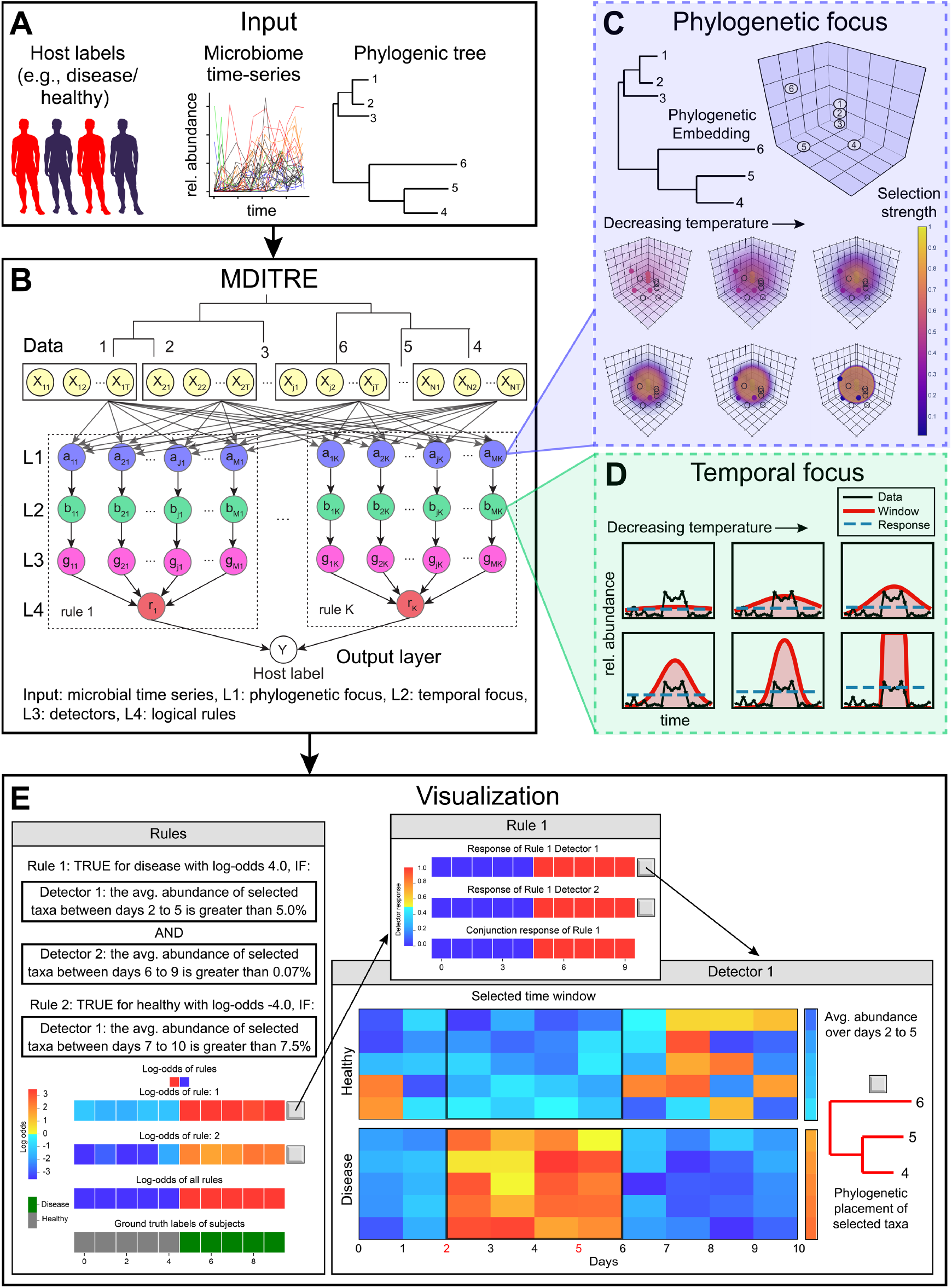
MDITRE efficiently learns rule-based machine learning models that predict host status from microbiome time-series data. **(A)** The inputs to MDITRE consist of binary labels for subjects (e.g., disease vs. healthy), relative abundances of microbiomes over time (derived from either 16S rRNA gene amplicon or metagenomics sequencing), and a phylogenetic tree. **(B)** MDITRE efficiently learns human-interpretable rules from the input data, using continuous relaxation techniques that render the model fully differentiable and amenable to deep-learning optimization techniques. The model consists of a five-layer neural network that includes specialized layers to learn phylogenetic and temporal features and to combine these features into rules. **(C)** Schematic of phylogenetic focus nodes, which embed taxa based on their phylogenetic distances and learn groups of related taxa to be used in rules. With decreasing temperature, the groups become sharper. **(D)** Schematic of temporal focus nodes, which learn relevant time-windows to be used in rules. With decreasing temperature, time-windows becomes sharper. **(E)** The output of MDITRE is a set of human-readable rules. Each rule consists of a conjunction (“AND”) of detectors that each selects a relevant set of taxa, a time-window and a threshold (for either aggregated abundance or rate-of-change of abundance). A graphical interface allows users to view per-subject strengths of rules (log-odds), and visualizations of detectors’ activations and components.

To create MDITRE, we introduced a set of relaxations, or continuous approximations, to discrete variables in the original MITRE model. These approximations render the MDITRE model fully differentiable, and thus amenable to highly parallel hardware-accelerated learning. Our original MITRE method effectively enumerated all subsets (phylogenetic subtrees) of microbes, time-windows, and abundance thresholds and then used a sampling-based inference algorithm to probabilistically explore this high-dimensional combinatoric space. MDITRE does away with explicit enumeration of features and instead directly (and continuously) parameterizes the model space. We accomplished this through several modeling innovations, including what we term microbiome or temporal *group focus* functions (**Figure 1C,D**), which perform “soft” selections over sets of microbes or time-points. To incorporate prior biological information into the microbe features, in terms of phylogenetic relationships, we introduced an embedding in phylogenetic space that “anchors” group focus functions. We also employed a relaxation of the logical AND operation, which was inspired by Neural Arithmetic Units^27^. As with our previous fully Bayesian model, we placed prior probability distributions on variables in the MDITRE model to incorporate biological information or to encourage model sparsity (i.e., total number of detectors or rules). We used relaxed versions of probability distributions to maintain differentiability. See Methods and Supplemental Methods for complete details.

### MDITRE was implemented in an open-source software package that uses standard deep learning libraries and has a graphical user interface

MDITRE can be represented as a five-layer Neural Network (**Figure 1B**), which allows us to directly leverage standard deep learning software packages. The top-most layer (layer 1) performs phylogenetic focus, generating outputs that are aggregated abundances of bacteria within the phylogenetically focused regions (**Figure 1C**). The temporal focus layer (layer 2) computes the average (or the rate of change) of its input (phylogenetically focused abundances) over temporally focused time-windows (**Figure 1D)**. The detector layer (layer 3) computes “soft” binary detector activations based on its inputs and detector thresholds. The rule layer (layer 4) performs “soft AND” operations over the input detector activations and then sends rule activations on to the last layer. Finally, the classification layer aggregates the rule activations from the previous layer to predict host labels.

We implemented MDITRE in Python using the PyTorch^28^ deep learning library, which fully supports GPU hardware acceleration, and have made the software package available under an open-source license. For model learning, we use standard gradient-descent based approaches in Pytorch, to perform maximum a posteriori estimation of model parameters. See Methods and Supplemental Methods for complete details. In addition to learning the model, the software provides a graphical user interface for visualizations of the learned rules, which allows end-users to readily interpret outputs (**Figure 1E**); we provide illustrative examples of these visualizations through use-case scenarios described below. We also provide a tutorial, which guides users step-by-step from dataset processing to final interpretations of outputs, to facilitate ease-of-use (see Methods).

### MDITRE performed comparably to our previous method MITRE but with up to orders of magnitude faster run-times

We first benchmarked MDITRE’s predictive performance on semi-synthetic time-series datasets generated using MITRE’s data simulation procedure. Briefly, simulated data was generated from real data using a parametric bootstrapping-type procedure, and perturbations of one or two randomly chosen microbial clades were added to subsets of artificial subjects to simulate diseased/dysbiotic states. Simulations were run for different numbers of artificial subjects ranging from 20 to 1024, and for different numbers of time-points ranging from 6 to 30 (for the case corresponding to 32 subjects). These ranges were chosen to correspond to sizes of real datasets and to test the scalability of the methods to larger dataset sizes. As with our previous comparisons^17^, we also benchmarked against L1 regularized logistic regression (L1) and random forest (RF), which are an interpretable linear method and a black-box nonlinear method, respectively. Our metric for comparison was five-fold cross-validated F1-scores (harmonic mean of precision and recall). Variability of performance was quantified using ten simulations of each synthetic dataset, and ten runs of each algorithm over each dataset.

For semi-synthetic data, MDITRE almost always performed comparably to MITRE and outperformed LR and RF (**Figure 2A,C,E,G**). For the case with one simulated perturbation and increasing numbers of subjects (**Figure 2A**), MDITRE performed comparably to MITRE on every case (*p*-values > 0.05; Mann-Whitney U Test, **Table S1**), excluding the simulations with 32 subjects, in which MITRE very slightly outperformed MDITRE (3% lower average performance for MDITRE). Moreover, MDITRE also achieved better performance compared to L1 and RF on every case (*p*-values < 0.05), excluding the simulations with 20 subjects (where L1 and RF performed comparably to MITRE) or 24 subjects (where L1 performed comparably to MITRE). For the case with two simulated perturbations and increasing numbers of subjects (**Figure 2C**), MDITRE performed comparably to MITRE (*p*-values > 0.05; **Table S2**) on all the cases. Moreover, MDITRE outperformed L1 and RF methods on all the cases (*p*-values < 0.05), except for the case with 32 subjects, where it performed comparably to RF (*p*-value 0.06). Overall, similar to what we previously reported^17^, for increasing numbers of subjects, we noticed a general increase in performance for all the methods, which eventually plateaued on cases with >48 subjects. For all cases with increasing numbers of time-points (**Figure 2E,G**), MDITRE achieved comparable performance to MITRE (*p*-values > 0.05; Table S3 and S4). MDITRE outperformed L1 and RF in most cases with increasing numbers of time-points, except for a few cases in which the methods performed comparably (**Table S3, S4**). Similar to what was reported for MITRE in^17^, for the cases with one simulated perturbation, we found no significant increase in performance with increasing time-points per subject, while for the case with two simulated perturbations, we found only a slight increase in performance with an increasing numbers of time-points. Both trends could be explained by the fact that sampling additional time-points that can lie outside the perturbation windows of interest does not provide any additional information to the model, while potentially adding noise that makes prediction more challenging.

**Figure 2:**
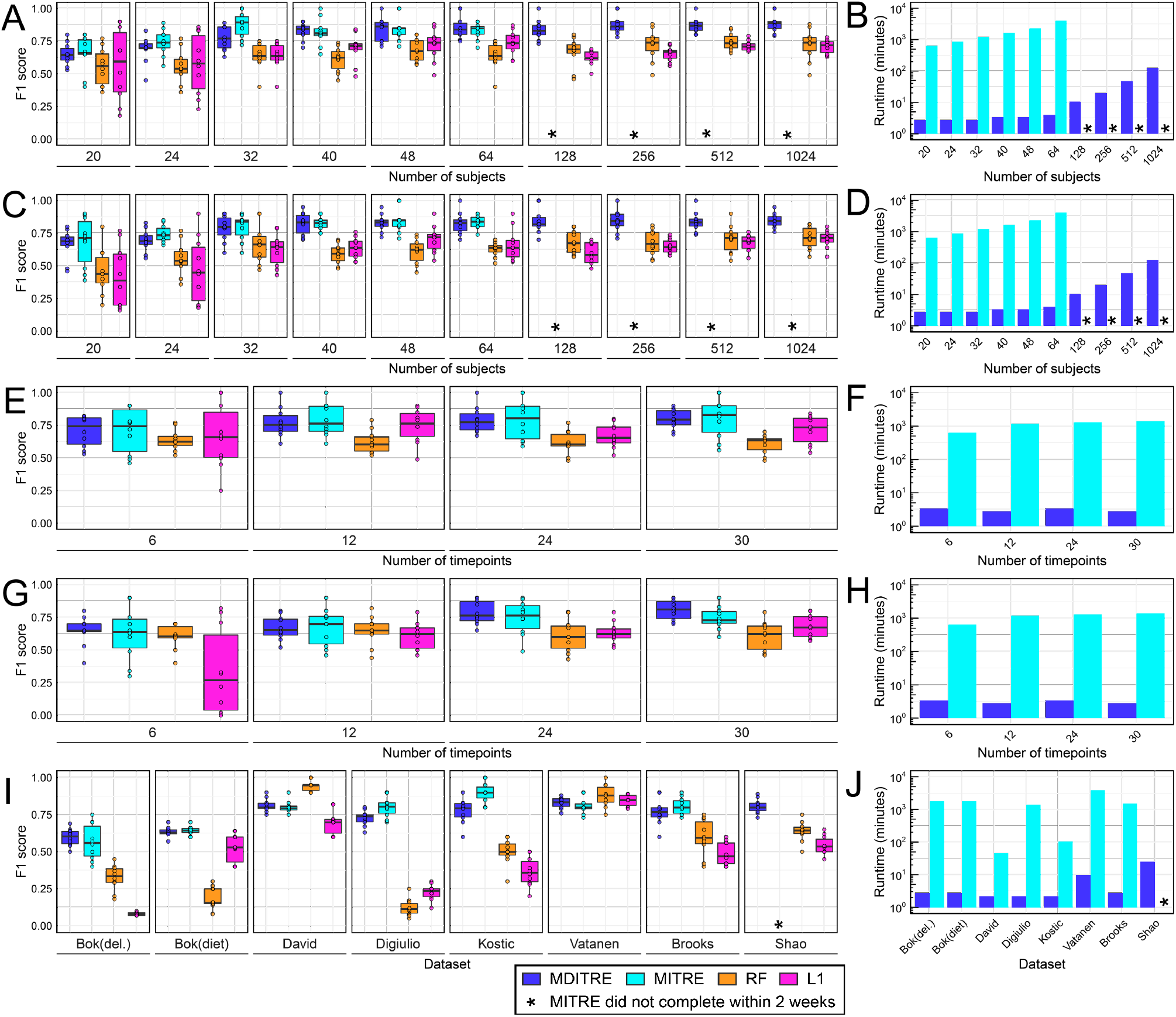
MDITRE achieved comparable performance to MITRE in almost all cases on both semisynthetic and real microbiome data, while running up to orders of magnitude faster. Classification performance for all methods was assessed via five-fold cross-validated F1 scores (harmonic mean of precision and recall), and variability was assessed with ten independent simulations or runs with different initializations. **(A,C,E,G)** show performance on semi-synthetic data for: **(A)** one or **(C)** two microbial clades perturbed with increasing numbers of subjects, **(E)** one or **(G)** two microbial clades perturbed with increasing numbers of timepoints and 32 subjects; **(B, D, F, H)** show corresponding runtimes (in log10 minutes). **(I)** Performance on real data, and **(J)** corresponding runtimes. RF = Random forest, L1 = L1 regularized logistic regression. Bok = Bokulick, del. = delivery.

MDITRE achieved significant runtime speedups over MITRE in all cases on semi-synthetic data (**Figure 2D,F,H, Table S6**), with particularly impressive speedups of >1000X on cases with larger numbers of subjects. Further, we were unable to complete benchmarking of MITRE for 128 subjects or greater, because in these cases MITRE still had not completed runs after two weeks of compute-time on our cluster; in contrast, MDITRE ran on the case with the largest number of subjects, 1024, in approximately 2 hours. This huge speedup in the runtime of MDITRE is attributable to its fully differentiable architecture that enabled us to implement an efficient gradient-descent-based learning algorithm, in contrast to the much slower MCMC-based learning algorithm employed by MITRE.

We next benchmarked MDITRE’s predictive performance on eight classification tasks from seven published human microbiome datasets: (1) Bokulich et al.^2^, a study of gut microbiomes of 37 infants sampled over the first two years of life, with two separate classifications tasks of diet (breastfed versus formula) and birth-mode (vaginal versus C-section), (2) David et al.^6^, a study of microbiomes of 20 healthy adults receiving dietary interventions (animal versus plant-based), (3) DiGiulio et al.^29^, a study of vaginal microbiomes of 37 pregnant women (at term versus pre-term delivery), (4) Vatanen et al.^30^, a study of gut microbiomes of 117 children sampled over the first three years of life (Russian versus Estonian/Finnish nationality), (5) Kostic et al.^4^, a study of gut microbiomes of 17 infants sampled over the first three years of life (normal versus development of type 1 diabetes), (6) Brooks et al.^31^, a study of gut microbiomes of 30 infants sampled over 75 days (vaginal versus C-section), and (7) Shao et al.^32^, a study of gut microbiomes of 282 infants (after filtering for subjects with fewer than three time-points) sampled over 424 days (vaginal versus C-section). Datasets 1-4 consisted of 16S rRNA amplicon sequencing data, and datasets 5-7 consisted of shotgun metagenomics data. See Methods for a complete description of bioinformatics and preprocessing of datasets. For benchmarking, we used the same metric as for the semi-synthetic data. Performance variability was computed over ten runs on each dataset with each method. For real data (**Figure 2I, Table S5**), MDITRE performed comparably to MITRE and outperformed the L1 and RF methods in almost all cases, while achieving massive speedups over MITRE. MDITRE had comparable performance to MITRE on six of the eight classification tasks (*p*-values > 0.05; Mann-Whitney U Test), while slightly underperforming MITRE on the DiGiulio et al. and Kostic et al. datasets (8.07% and 11.67% lower average performance, respectively). MDITRE significantly outperformed the L1 or RF methods on all the datasets, except for the Vatanen et al. dataset (comparable performance to L1 and RF) and the David et al. dataset (16% lower average performance for both MDITRE and MITRE, relative to RF; all methods outperformed L1). In terms of runtimes (**Figure 2J, Table S6**), MDITRE achieved speedups over MITRE ranging from 86-1150X. MITRE was unable to run on Shao et al., due to the size of the dataset, whereas MDITRE completed analysis of this dataset in 24 minutes. We note that for the three cases in which MDITRE underperformed other methods, these were the most imbalanced or small datasets. DiGiulio et al. is highly imbalanced with only 6 of 37 subjects belonging to the “positive” group. Kostic et al. is the smallest datset with 17 subjects, and David is the second smallest with 20 subjects. Thus, predictive accuracy results on these datasets should be interpreted with caution, as high sample imbalance or small sample size could lead to less reliable estimates for cross-validated F1 scores. Overall, our results on both semi-synthetic and real datasets demonstrate that MDITRE, which uses continuous relaxations to approximate discrete variables in our MITRE model, achieves competitive classification performance to our original method, while running orders of magnitude faster and scaling to much larger datasets.

### MDITRE discovered human interpretable rules that automatically focused on biologically relevant taxa and time windows

We next examined the interpretability of MDITRE’s outputs, which we demonstrate through two case studies. In the prior section, we focused on the objective measure of predictive performance. However, for many microbiome applications, the critical tasks are discovering relationships between the microbiome and the host or finding clinically useful biomarkers, rather than pure prediction. For these purposes, model interpretability, which is inherently domain-specific and subjective^23^, is the key property of interest. Through the case studies in this section, we illustrate powerful features of the rules and visualizations that MDITRE returns, which facilitate interpretability tailored to the domain of microbiome time-series analyses through: (1) automatic focus on relevant groups of taxa or single taxa that differentiate hosts, (2) automatic focus on time windows in which the microbiome is differentially changing depending on host status and, (3) human-readable rules with “AND” and “OR” logic, which can capture rich patterns of dynamics or host variation, while remaining easily understandable.

#### Case study one: the relationship between diet and the microbiome in infants

Our first case study used the dataset of Bokulich et al., which analyzed gut microbiomes of 37 infants during the first two years of life, using 16S rRNA amplicon sequencing. Note that although there were samples over two years, they trailed off significantly after the first year of the study, and our preprocessing procedure to ensure sufficient samples in time-windows truncated the analyzable data to the first 375 days (see Methods and^17^ for complete details). Given the task of classifying infants as receiving either breast milk or formula predominant diets, MDITRE learned two rules. **Figure 3** illustrates information available through MDITRE’s graphical user interface, which aids in interpreting these rules. **Figure 3A** illustrates how the rules are combined in a logical “OR” to predict host labels. In this case, we can see that each rule alone can correctly classify most infants, but not necessarily with high odds. However, when the rules are combined, the separation between the classes is improved (i.e., higher odds of classifying as either formula or breastfed), and several infants that would be misclassified by individual rules were correctly classified by the combined rules.

**Figure 3:**
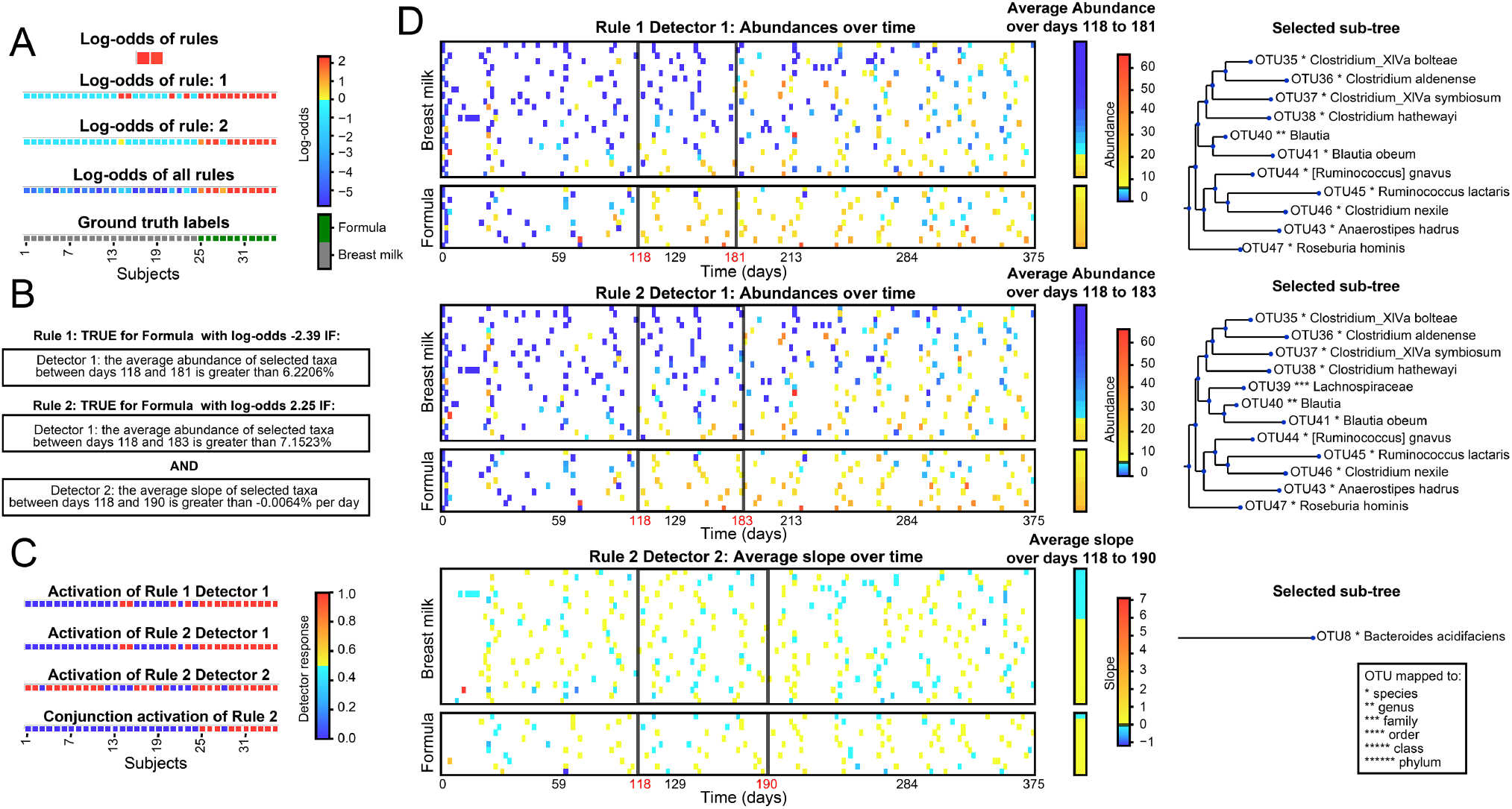
Case study one, MDITRE rules that distinguish predominantly breast versus formula-fed infants based on gut microbiome changes over time. Visualizations output by the MDITRE graphical user interface are shown, for analysis of data from Bokulich et al., which tracked gut microbiomes of 37 infants using 16S rRNA gene amplicon sequencing. **(A)** MDITRE identified two rules that in combination accurately differentiate infants who were predominantly breast versus formula-fed. The contribution of each rule, in terms of log-odds, is shown for each subject. **(B)** Human-readable logic of the rule. **(C)** Per-subject activations (“truth” values) for each detector in each rule, and the conjunction (combined effects) of detectors for rules that have more than one detector. **(D)** Visualization of the per-subject data aggregated by each detector, as well as the detector’s window-time (black boxes), threshold for either abundance or slope (black line on abundance or slope legend) and selected taxa.

By clicking on a rule, the user can then see an English description (**Figure 3B**) and a visualization of how the rule’s detectors combine in a logical “AND” to determine the rule’s final truth value (**Figure 3C**). For brevity, we discuss the second rule, which contains two detectors. The first detector is true for all the predominantly formula-fed infants, but also four of the breast-fed infants. In contrast, the second detector is true for more infants overall (both formula- and breast-fed) but is false for the four breast-fed infants the first detector was true for; this suggests the second rule finds complementary information to aid in defining breast-fed status for a group of infants. The conjunction of the detectors then provides a final rule that is true for all but one formula-fed infant, and false for all the breastfed infants.

By clicking on the detectors, the user can then view visualizations (**Figure 3D**) of the time windows and taxa selected by each detector. Both detectors for the second rule focus on approximately the same time window, between about four to six months. Interestingly, this is a period directly proceeding the introduction of the first solid foods for most infants (and which may occur earlier for formula-fed infants^33^, thus explaining why MDITRE may have selected this time-period rather than a later one). Because infant microbiomes are extremely variable for the first few months of life^34^, the automatically selected time window seems to reflect an optimal period for differentiating formula-versus breastfed infants: when the microbiome has had time to equilibrate, but before a new perturbation introduced by solid foods.

The first detector for the second rule, an aggregate abundance type, selected twelve taxa in the Order Clostridiales, including *Clostridium, Blautia, Ruminococcus, Anaerostipes*, and *Roseburia* genera. This detector, which is true when the aggregate abundance of these taxa is greater than ~7%, detects all the formula-fed infants and four of the breast-fed infants (because the study only reported the predominant feeding mode, it is possible these infants also received relatively more formula than others in the breastfed group). By focusing on the aggregated abundance of these taxa, the detector has automatically uncovered a group of phylogenetically related microbes that may not all be present in one individual or all in high amounts, but in aggregate may reflect a common biological function. Indeed, many of the selected taxa are strict anaerobes that metabolize more complex nutrient sources, including starches and lipids^35^ that may be present in formula. Interestingly, the visualization produced by MDITRE suggests that after the selected time window, the abundance of these taxa became increasingly difficult to distinguish between formula- or breast-fed infants, which may reflect similar diets post-liquid-foods in both groups. This again highlights the ability of MDITRE to automatically focus on relevant time windows.

The second detector, a slope-type, selects a single taxon, *Bacteroides acidifaciens*, and is true if this taxon is increasing. This detector was not only true for all but one formula-fed infant but also many breast-fed infants; however, it was false for the four breastfed infants identified by the first detector. *Bacteroides acidifaciens* has been shown to increase with higher fiber diets^36^. Thus, this detector may be capturing a gradient of the introduction of solid foods in the infants, with delayed introduction of solid foods for some breastfed infants.

#### Case study 2: temporal series of changes in the microbiome preceding onset of type 1 diabetes

Our second case study used the Kostic et al dataset, which tracked children’s gut microbiomes over the first three years of life using shotgun metagenomics, and also assessed the onset of type 1 diabetes (T1D). In this case, MDITRE found a single rule (**Figure 4A-B**), which contains three detectors of slope-types (**Figure 4C**), covering progressive time windows throughout the study: approximately 5 to 15 months, 13 to 22 months, and 17 to 26 months (**Figure 4D**). In all cases, each detector was true for increases of the selected taxa, which occurred in children who did not develop T1D. The taxa selected were *Escherichia coli* for the earliest detector, *Streptococcus* and *Coprobacillus* for the middle detector, and *Faecalibacterium prausnitzii* for the latest detector (**Figure 4D**). Interestingly, these taxa are progressively anaerobic and specialized to the gut, consistent with community succession events that occur in the normal developing infant gut^37^. Thus, the rule MDITRE discovered to classify T1D versus non-T1D infants specifies a temporal pattern of events that appears to detect normal microbiome succession events in healthy infants, which are absent or blunted in infants who later develop T1D.

**Figure 4:**
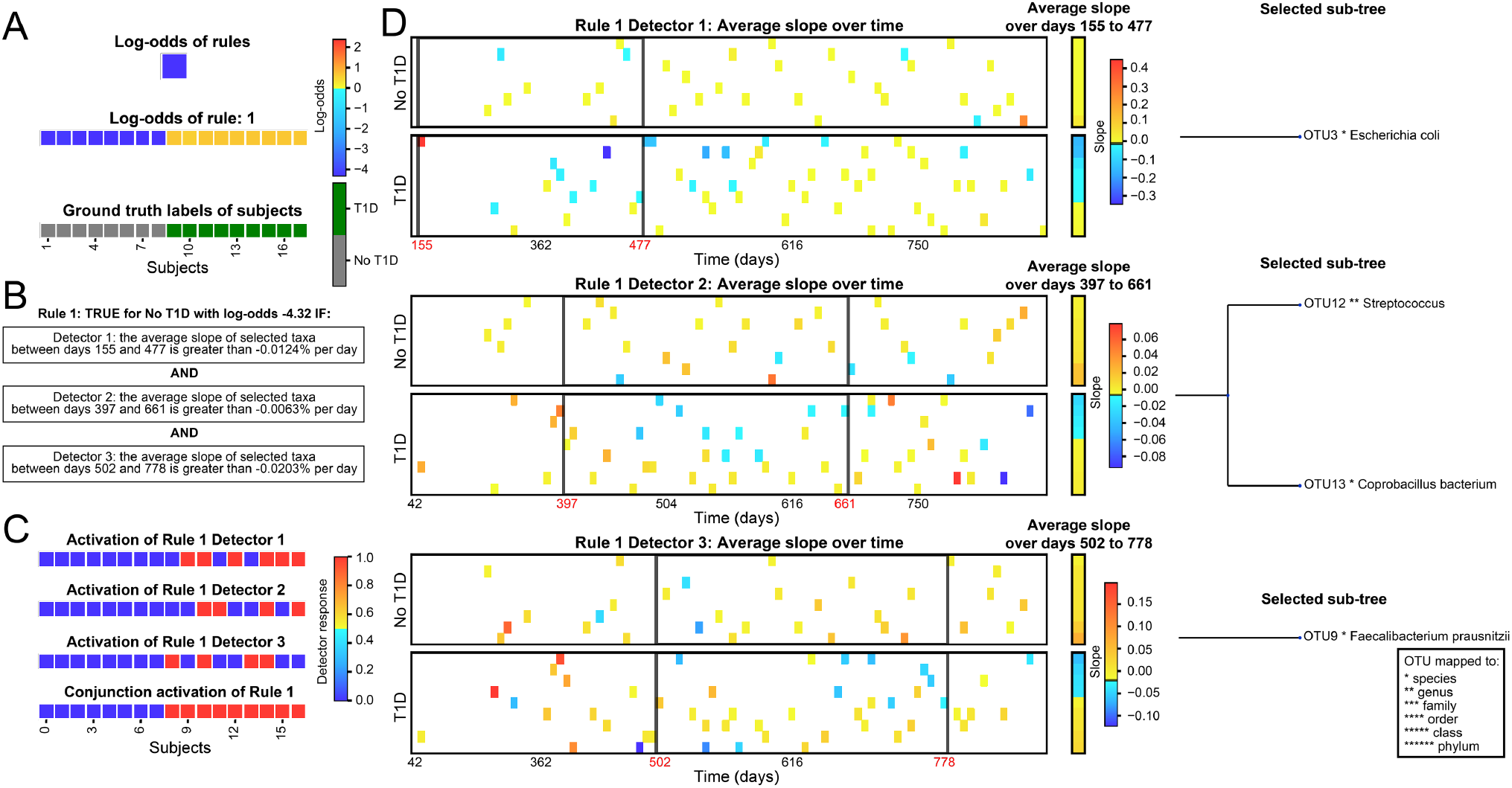
Case study two, MDITRE rules that predict children who developed type 1 diabetes versus those who did not, based on a series of gut microbiome changes over time that precede disease onset. Visualizations output by the MDITRE graphical user interface are shown, for analysis of data from Kostic et al., which tracked gut microbiomes of 17 infants during the first three years of life using shotgun metagenomics sequencing. **(A)** MDITRE identified one rule that predicts which children developed type 1 diabetes. **(B)** Human-readable logic of the rule, which has three detectors. **(C)** Per-subject activations (“truth” values) for each detector and the conjunction (combined effects) of the detectors. **(D)** Visualization of the per-subject data aggregated by each detector, as well as the detector’s window-time (black boxes), threshold for slope (black line on legend) and selected taxa. Note that the detectors identify a pattern of temporal changes that appears to correspond to normal microbiome succession events that are absent or blunted in infants who later develop type 1 diabetes.

## Discussion

We have presented MDITRE, software for learning human-interpretable models that predict host status from microbiome time-series data, which achieves comparable predictive performance to our original method in almost all cases, while running up to orders of magnitude faster. Moreover, we have introduced new visualization capabilities and shown through case studies that our method uncovers rich but readily interpretable and biologically relevant patterns in microbiome datasets. To achieve these improvements, we introduced several innovations, including relaxation techniques that operate on temporal or phylogenetic information to render models fully differentiable. With these innovations, we took advantage of standard machine learning libraries that support GPU acceleration and are easily deployable on different operating systems.

We foresee several directions for future work. MDITRE currently supports binary host labels, which we found to be the most common scenario for microbiome datasets. However, the model could readily be extended to multiclass learning. In addition, the model could be extended to time-to-event (survival) prediction tasks, which are relevant for some recent microbiome studies, such as predicting the risk of nosocomial infections^38,39^ and the likelihood of disease resolution^19^. Similarly, longitudinal microbiome studies are beginning to include multiple data modalities^40^. Due to its layered nature, MDITRE can be extended to incorporate additional multi-omics data, such as metabolomics or transcriptomics information. Finally, although MDITRE is a Bayesian model, we used a maximum a posteriori inference procedure that cannot estimate uncertainty throughout the model. Future work could involve applying inference techniques, such as Variational Inference^41^ or Hamiltonian Monte-Carlo^42^, which take advantage of MDITRE’s differentiability while also estimating the posterior distribution.

Overall, we have introduced MDITRE, a new software package that improves on our prior work with orders of magnitude faster run-times and expanded visualization capabilities, to address an important gap in the field: linking changes in the microbiome over time to the status of the host. Benchmarking on semi-synthetic and real datasets shows that our software performs on par with or outperforms a high-capacity interpretable machine learning method (random forests) in almost all cases, in terms of predictive performance, while returning human-interpretable rules that capture domain-specific features of microbiome time-series data. We believe that MDITRE will provide a valuable tool for the microbiome research community, fostering new insights into how changes in the microbiome over time maintain health or lead to disease in humans and other species.

## Online Methods

### MDITRE model

We describe the model in terms of a five-layer Bayesian neural-type architecture: (1) phylogenetic focus, (2) temporal focus, (3) detectors, (4) rules, and (5) classification. Supplemental Methods provides an alternate view, as a graphical (plate) model showing the probabilistic structure of MDITRE. The model is fully differentiable, and we implemented maximum a posteriori (MAP) inference using gradient-descent.

#### Phylogenetic focus layer

This layer learns localized features by aggregating abundances of phylogenetically related OTUs. Let P ∈ ℛ^*𝒩*×*𝒩*^ denote a dissimilarity matrix, e.g., pairwise phylogenetic distances. To allow differentiability, we embed *p* into a *D*-dimensional space using Principal Coordinate analysis. Our software automatically chooses the value of *D* for each dataset by running a two-sample Kolmogorov-Smirnov (KS)-test using the original phylogenetic distances and the distances after embedding, for *D* values ranging from 1 to 30, and choosing the lowest value of *D* with a *p*-value > 0.05 (the lowest number of dimensions such that the original and embedded distributions of distances are not significantly different). Let E ∈ ℛ^*𝒩*×*𝒟*^ denote the resulting embedding matrix. We assume that each detector *j* in rule *k* has a phylogenetic “center” γ_kj_ ∈ ℛ^𝒟^ and scalar radius κ_kj_; we assume Normal and Lognormal priors on γ_kj_ and κ_kj_ respectively.

The distance *ξ*_*kji*_ between detector *j*’s phylogenetic center (in rule *k*) and OTU *i′s* phylogenetic embedding is defined as:

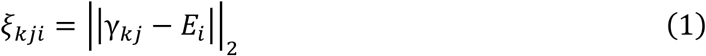

The layer’s output a_*skj*_, for each host *s*, is then a “soft” aggregation of OTUs that fall within detectors’ phylogenetic centers:

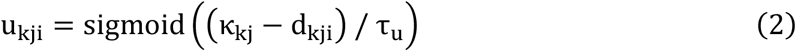

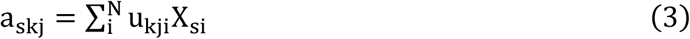

Here, *X*_*sit*_ denotes a microbiome measurement for time-series *s* at time *t* for OTU *i* of an *N*-dimensional data source (e.g., relative abundances of OTUs from 16S rRNA amplicon or metagenomic shotgun sequencing). The parameter *τ*_u_ is a temperature parameter that increases the sharpness of focus with lower temperatures.

We place priors on the phylogenetic centers and radii to encourage focus on interpretable regions of the tree. For κ, we use a Normal prior with mean set to the median of all family-level distances of OTUs and variance set to the median of all variances of family-level distances of OTUs, both of which are derived from the embedding of the reference phylogenetic tree. See Supplemental Methods for complete details.

#### Temporal focus layer

This layer models two types of localized features in the temporal space: (1) average abundances over a time-window, or (2) rates of change (slope) of abundances over a time-window.

##### Average abundances over time-windows

we use a similar approach to that described for the phylogenetic focus layer. Each detector probabilistically selects a time-window center and extent, and then data from time-points in that window are “softly” averaged. Let μ_kj_ denote the time-window center for detector *j* in rule *k* and σ_kj_ its corresponding extent (length). We model the random variables μ_kj_ and σ_kj_ in terms of the fraction of the total experiment length; see Supplemental Methods for complete details.

The output of this layer, *b*_*skj*_, for each host *s*, is then a (soft) average over the phylogenetically focused microbial abundance data from the previous layer:

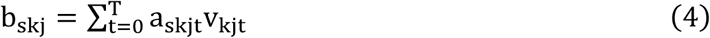

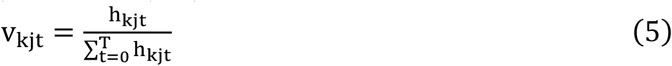

Here, *h*_*kjt*_ are (soft) indicators as to whether time-points occur in the time-window. We compute *h*_*kjt*_ as follows, using a relaxed approximation to the Heaviside boxcar function with temperature τ_v_.

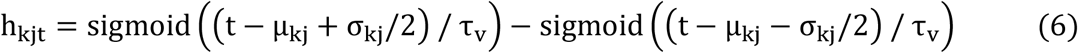

##### Rate of change of abundances (slopes) over time-windows

we use a similar approach as for average abundances over time-windows, described above. Briefly, we (softly) estimate the slope over the time-window using weighted Ordinary Least Squares (OLS) regression with weights 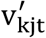 computed as for average abundances. See Supplemental Methods for complete details.

We set priors on random variables in the temporal focus layer to encourage time-windows that correspond to intervals of time relevant to the underlying studies’ experimental designs. These settings encode prior beliefs that time-windows encompass 30% of the total study duration and are centered at the mid-point of the study. However, we intend for these priors to be relatively weak, and thus set large variances to create diffuse priors.

#### Detector layer

The detector layer takes its inputs from the previous layers, which consist of phylogenetically and temporally focused features, and computes activations according to whether the feature is above learned thresholds. By design, the detectors effectively form human-interpretable clauses, i.e., for detector *j* in rule *k*, “*TRUE if b*_*skj*_ *(the [aggregated abundance / rate of change of abundance] of organisms within the detectors’ phylogenetic radius and time window) is above threshold η*_*kj*_.” To maintain differentiability, we model the activations using sigmoidal responses with a temperature parameter that is annealed toward increasing sharpness of detectors; we place uniform priors on the threshold random variables. We set the maximum number of detectors per rule to ten, based on our prior work^17^, where we showed this to be a very liberal setting for microbiome datasets (e.g., most rules contained only a few detectors).

#### Rule layer

The rule layer takes the detector activations as inputs and performs a relaxed logical conjugation. We use a relaxation inspired by arithmetic-operation learning networks, which has less impact from vanishing gradients^27^ than an exponentiation-based relaxation. The activation for rule *k* for time-series *s* is thus:

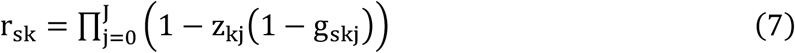

Here, *J* is the maximum number of detectors per rule. The random variables z_kj_ (softly) select which detectors are relevant to each rule, using a sigmoid function applied to Normally distributed latent variables; see Supplemental Methods for complete details.

To encourage model parsimony, we place Negative Binomial priors on the number of detectors per rule 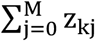, parameterized by mean θ_z_ and variance 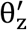:

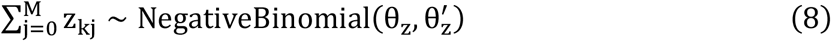

For the datasets analyzed, we set θ_z_ = 1 and 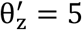, encoding a prior that the 98th percentile of this distribution equates to 50% of model capacity (≤5 active detectors per rule). We set the maximum number of rules to ten, based on our prior work^17^, where we showed this to be a very liberal setting for microbiome datasets (e.g., most predictors had fewer than three rules).

#### Classification layer

The classification layer takes the rules’ activations and combines them via a logistic regression model to predict the binary label for each subject. We place diffuse priors on the regression coefficients. Analogous to the detector layer, we introduce random variables *q*_*k*_ that (softly) select which rules are relevant to the final predictor; we similarly place a Negative Binomial prior on the total number of rules in the model, to encourage parsimony. As with the detector layer, we set the mean of this prior to 1 and the variance to 5. See Supplemental Methods for complete details.

### Initialization

To initialize the phylogenetic focus layer centers and radii, we performed *K*-means clustering on the OTUs in the phylogenetic embedding space (with *K* equal to the maximum number of detectors per rule, *J*), and then set the initial detector phylogenetic centers and radii from the *K*-means output. To initialize the temporal focus layer time-window centers and durations, we set initial values to randomly selected time windows from *N*_*w*_ consecutive segments of the total experiment duration. We compute *N*_*w*_ to be the maximum number of consecutive time intervals while ensuring the presence of at least 2 samples per subject in each interval. The procedure is explained in the algorithm below:

#### Algorithm 1: Initialize detector time-windows and centers

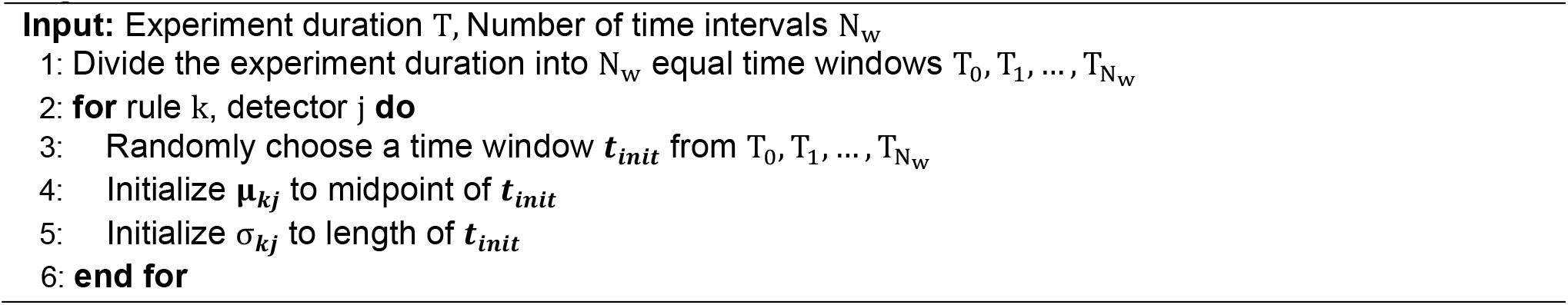

To initialize the detector layer abundance or slope thresholds, we set the initial values equal to the mean over all subjects of the aggregated abundances (or slopes) computed based on the initializations of the phylogenetic or temporal focus parameters as described above. We initialized logistic regression coefficients from a standard Normal distribution and set initial bias terms to zero. We initialized the rule and detector selectors to 0.5, which ensures equal probability of a rule or detector being active at the start of the training.

### Model training and testing

We performed MAP inference using RMSProp. We used a learning rate of 0.01 for the temporal focus layer parameters (μ, σ), and 0.001 for all the other parameters. We found that in our experiments, the temporal focus parameters needed a higher learning rate based on their scale, which is typically an order of magnitude higher than all the other parameters. For example, the temporal focus parameters typically are on the order of 10 – 100 days, whereas other parameters such as rule and detector selectors, thresholds and phylogenetic windows are typically on the order of 0 to 1. For full learning rate settings please refer to our code available here: https://github.com/gerberlab/mditre.

Temperature parameters were linearly annealed throughout learning toward sharpness. The temperature parameters for phylogenetic, temporal, rule and classification layers were annealed from 1 to 0.1. The temperatures for thresholds for detectors were annealed from 10^−2^ to 10^−3^ and from 10^−3^ to 10^−4^, for aggregated abundances and slopes respectively; these ranges correspond to the scales of abundances or slopes in data. The model was trained using RMSprop optimizer (with default parameters given here: https://pytorch.org/docs/stable/generated/torch.optim.RMSprop.html#torch.optim.RMSprop) for 2000 iterations on all the datasets, which was adequate to ensure convergence.

We used the following hardware configuration for benchmarking: Intel Xeon Silver 4116 CPU (2.1 GHz) with 24 cores and 48GB RAM and an NVIDIA Tesla V100 GPU.

### Details on code, implementation, and availability

The model was implemented in Python 3.6 and Pytorch 1.6^28^. The NumPy 1.21, Scikit-learn 0.24, Matplotlib 3.4, SciPy 1.6 and ETE3 3.1 libraries were also used.

The full source code and documentation is available here: https://github.com/gerberlab/mditre under a GPL 3.0 license.

Scripts to reproduce all results in the manuscript are available at: https://github.com/gerberlab/mditre/tree/master/mditre_paper_results

We also provide a tutorial here: https://github.com/gerberlab/mditre#usage that describes how to perform tasks such as data loading and preprocessing, running the model, and exploring the output through the graphical user interface.

### Datasets and Bioinformatics

The datasets corresponding to Bokulich et al, David et al, Vatanen et al, Digiulio et al, and Kostic et al. were all downloaded as Python pickle objects from: https://github.com/gerberlab/mitre/tree/master/mitre. They were input into MDITRE without any further preprocessing.

The datasets for Brooks et al. and Shao et al. were first downloaded from the R Bioconductor Package: (https://bioconductor.org/packages/release/data/experiment/html/curatedMetagenomicData.html) and then transformed into a Python pickle object and input to MDITRE for modeling. A 5% prevalence cutoff was used to filter taxa to include in the modeling for these two datasets.

The phylogenetic prior for 16s rRNA datasets was calculated using a reference phylogenetic tree with 7500 OTUs available from: https://github.com/gerberlab/MDSINE2_Paper/blob/master/analysis/files/phylogenetic_placement_OTUs/phylogenetic_tree_full.nhx. Placements of OTUs on the tree was performed by running the software pplacer (https://matsen.fhcrc.org/pplacer/). For the metagenomics datasets phylogenetic distances between taxa were determined after mapping the taxa to a reference tree of 9700 strains available within the Metaphlan2 package: (https://github.com/biobakery/MetaPhlAn/blob/master/metaphlan/utils/mpa_v30_CHOCOPhlAn_201901_species_tree.nwk).

## Acknowledgements

We acknowledge the use of the computational resources of the Center for Scientific Computing and Visualization Research at UMass Dartmouth and from the Mass Green High-Performance Computing at UMass Medical School. In particular, the CARNIE cluster which was funded by ONR DURIP Grant No. N00014181255. We acknowledge guidance from Richard Croswell for running the MITRE software.

## MDITRE Supplementary Methods

### 1 Probabilistic Model

In this section we describe the mathematical details of the MDITRE probabilistic model. Figure 1 provides a graphical model depiction. As described in the main text, the MDITRE model can also be viewed as a five-layer neural-type architecture. Each layer learns relevant features that form the interpretable rules, which are combined in the final layer to predict the class label. The layers are as follows:

- **Phylogenetic focus layer:** Learns relevant regions in an embedded space, and outputs aggregate abundances of phylogenetically similar OTUs.
- **Temporal focus layer:** Learns relevant time-windows, and outputs average abundances or rates of change over selected time-windows.
- **Detector response layer:** Learns thresholds (for abundances or rates of change of abundances) to and outputs (soft) truth values of clauses.
- **Rule response layer:** Learns which clauses (detectors) are relevant and outputs (soft) logical conjunctions of detector outputs.
- **Class prediction layer:** Learns which rules are relevant and how to weight relevant rules, and outputs class label predictions.

#### 1.1 Phylogenetic Focus Layer

This layer learns localized features by aggregating abundances of phylogenetically similar Operational Taxonomic Units (OTUs). Let *X*_*sit*_ denote a microbiome measurement from subject *s* at time *t* for OTU *i* of an *N*-dimensional data source (e.g., relative abundances of OTUs from 16S rRNA amplicon or metagenomic shotgun sequencing). Let *P* ∈ ℝ^*N×N*^ denote a phylogenetic distance matrix. We embed *P* into a latent space of dimensionality *D* (e.g., via PCoA on *P*) and denote the resulting embedding matrix as *E* ∈ ℝ^*N×D*^. We associate with each rule *k*, detector centers *γ*_*kj*_ ∈ ℝ^*D*^ and scalar radii *κ*_*kj*_, where *j* ∈ {1, …, *J*} (*M* detectors per rule).

**Figure 1:**
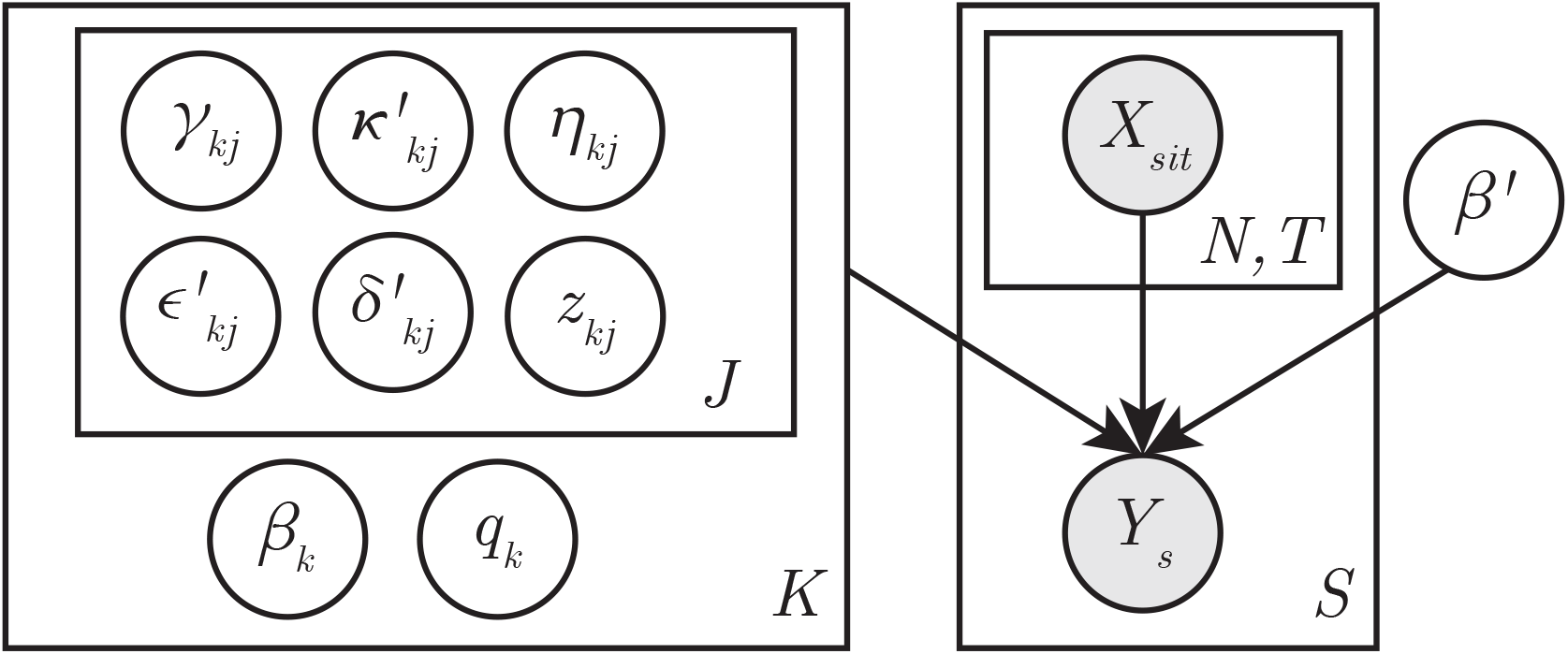
Graphical model depiction of the MDITRE probabilistic model. Observed variables, *X*_*sit*_ (microbiome time-series) and *Y*_*s*_ (class labels) are shaded. The nested plates on the left contain latent variables that model *K* rules and their associated detectors (*M* detectors per rule). The class label *Y*_*s*_ is predicted via logistic regression on the rules with weights *β*. The variables *z*_*kj*_ and *q*_*k*_ select which detectors or rules are active, respectively. Latent variables have prior probability distributions (not shown; see text for details).

We define the distance *d*_*kji*_ between the detector center *j* in rule *k* and OUT *i*’s phylogenetic embedding as:

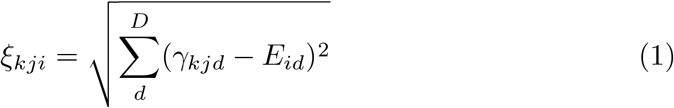

OTUs with distances within radius *κ*_*kj*_ are then softly selected for inclusion:

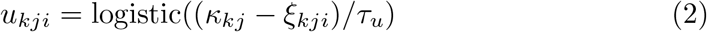

Here, *τ*_*u*_ is a temperature parameter that determines the selection sharpness.

The outputs of the layer are aggregated abundances, *a*_*skj*_, of softly selected OTUs, for each detector *j* in rule *k* in subject *s*:

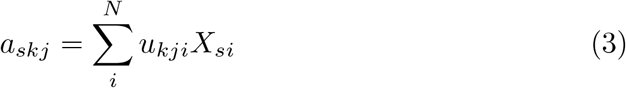

We place Normal priors on the detector centers *γ*_*kj*_:

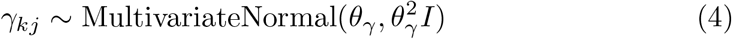

We place Lognormal priors on the detector radii *κ*_*kj*_:

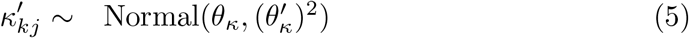

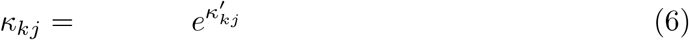

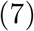

#### 1.2 Temporal Focus Layer

This layer learns two types of temporally localized features: (1) average abundances *b*_*skj*_ over time-windows, and (2) average rates of change of abundances (slopes) 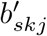 over time-windows.

Relevant time-windows are learned using a temporal focus mechanism that (softly) selects time-points. Each detector *j* in rule *k* learns a window center *μ*_*kj*_ and duration *σ*_*kj*_. We parameterize *σ*_*kj*_ in terms of fractions *ϵ*_*kj*_ of the total experimental duration *T* :

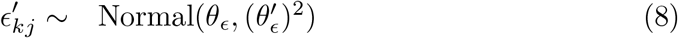

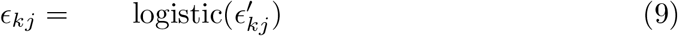

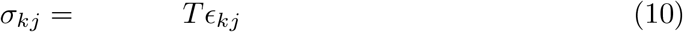

Note that the prior on 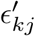 specified in Equation 9 can be used to encode knowledge or beliefs about relevant time-window durations.

We parameterize the window centers *μ*_*kj*_ similarly:

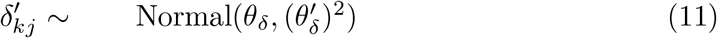

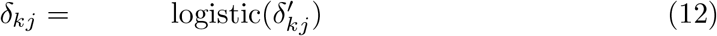

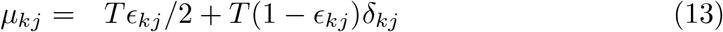

We use a relaxed boxcar function to compute (soft) selections *v*_*kjt*_, for each time-point *t*:

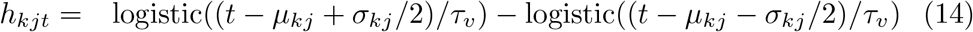

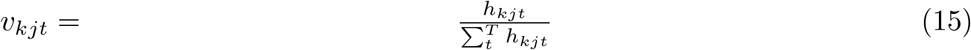

Here, *τ*_*v*_ is a temperature parameter controlling the sharpness of the approximation to the boxcar function.

##### 1.2.1 Average abundances over time-windows

The output is a (soft) average over the selected time-windows of abundances *a*_*skjt*_ (from the previous layer):

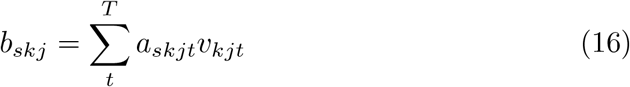

##### 1.2.2 Average rates of change of abundances (slopes) over time-window

The output is a (soft) slope over the selected time-windows of abundances *a*_*skjt*_ (from the previous layer):

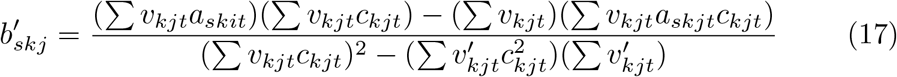

Here, *c*_*kjt*_ = *t* − *μ*_*kj*_. Note that Equations 17 is the solution to the Weighted Linear Regression problem with weights *v*_*kjt*_.

#### 1.3 Detector Activation Layer

This layer learns activation thresholds *η*_*kj*_ and 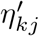 for each detector *j* in rule *k*, i.e., if the input (the phylogenetically and temporally focused abundance *b*_*skj*_ [or slope 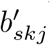]) is greater than *η*_*kj*_ (or 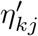, the detector is (softly) true. We place a Uniform prior on *η*_*kj*_ and 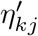 :

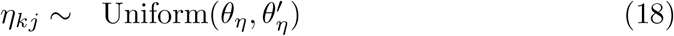

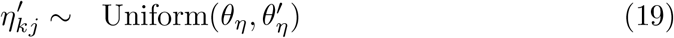

The output of the layer is given by:

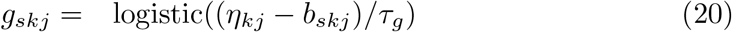

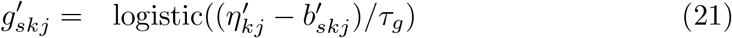

Here, *τ*_*g*_ is a temperature parameter that controls the sharpness of the activation.

#### 1.4 Rule Activation Layer

This layer learns which detectors are relevant and then outputs a (soft) logical conjunction of the selected detectors. We model the joint probability of detector selectors as:

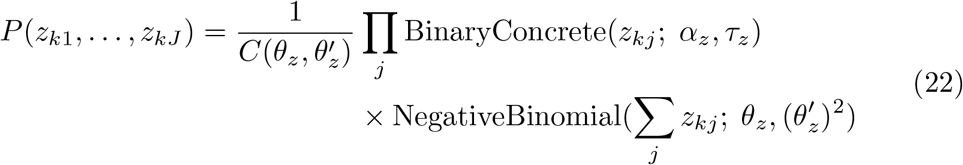

Here, each *z*_*kj*_ ∈ (0, 1) and *τ*_*z*_ is a temperature parameter for the BinaryConcrete distribution, which is a continuous relaxation of the discrete Bernoulli distribution. Note that we do not need to compute the normalization constant, 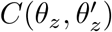, because *θ*_*z*_ and 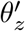 are fixed parameters that we are not optimizing over.

We use the following PDF of the BinaryConcrete parameterized by location *α*_*z*_ ∈ (0, ∞) and temperature *τ*_*z*_:

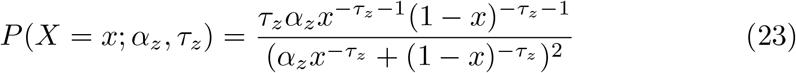

Equation 22 also places a prior probability distribution on the total number of active detectors in each rule through the NegativeBinomial distribution on the sum of detector selector variables. We use this distribution to encode model sparsity (inductive bias toward small numbers of detectors).

We use the following PDF for the Negative Binomial, which is parameterized in terms of its mean *θ*_*z*_ and variance *θ*_*z′*_ and allows for non-integer values:

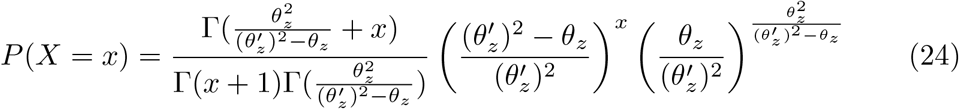

The outputs of the layer are rule activations, *r*_*sk*_, which are relaxed logical conjunctions (modeled as products) of selected detector responses *g*_*skj*_:

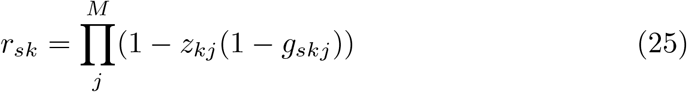

#### 1.5 Prediction Layer

This layer learns which rules are relevant and weights for the rules, which are then used to predict the probabilities of the binary class labels *Y*_*s*_. We model the (soft) rule selectors, *q*, analogously to the detector selectors described above:

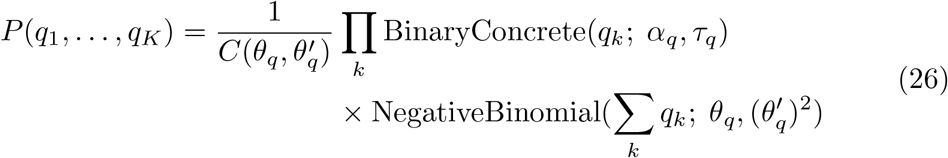

Analogous to the prior on detector selectors, Equation 26 places BinaryConcrete priors on the rule selectors and a NegativeBinomial distribution on the total number of active rules in the model. We use the latter to encode model sparsity (inductive bias toward a small number of rules).

We use a logistic regression model over the selected rules to predict class probabilities:

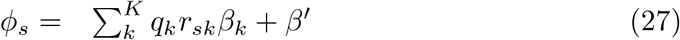

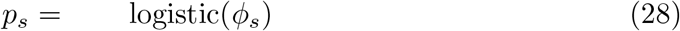

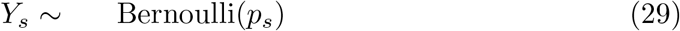

Finally, we place Normal priors on the regression coefficients (weights) *β*:

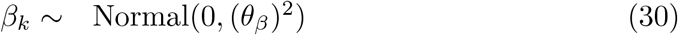

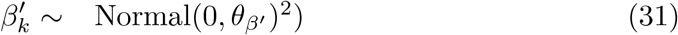

